# Variants of intrinsic disorder: structural characterization

**DOI:** 10.1101/721308

**Authors:** Sergio Forcelloni, Antonio Deiana, Andrea Giansanti

## Abstract

In a recent study, we have introduced an operational classification of the human proteome in three variants of disorder: ordered proteins (ORDPs), structured proteins with intrinsically disordered protein regions (IDPRs), intrinsically disordered proteins (IDPs). That classification was useful in functionally separating IDPRs from IDPs, which up until now have been generally considered as a whole. In this study, we corroborate this distinction by considering different physical-chemical and structural properties. Both ORDPs and IDPRs are enriched in order-promoting amino acids, whereas only IDPs show an enrichment in disordered-promoting amino acids. Consistently, ORDPs and IDPRs are preferentially located in the ordered phase of the charge-hydropathy plot, whereas IDPs are widespread over the disordered phase. We introduce the *mean packing - mean pairwise energy (MP-MPE) plane* to structurally characterize these variants even in the absence of a structural model. As expected for well-packed proteins, a negative linear correlation is observed between MP and MPE for ORDPs and IDPRs, whereas IDPs break this linear dependence. Finally, we find that IDPs have a more extended conformation as measured by the scaling law between the radius of gyration and the length of these proteins, and accordingly they have higher solubility and accessible surface area than ORDPs and IDPRs. Overall, our results confirm the relevance of our operational separation of IDPRs from IDPs and provide further validation of our criteria to separate IDPs from the rest of human proteome.

## 1. Introduction

Over the last two decades, the concept of *‘intrinsic disorder’* has emerged as a prominent and influential topic in protein science [1-7]. Intrinsically disordered proteins (IDPs) possess regions of primary sequence that are highly flexible in solution and lack a well-defined three-dimensional structure. Their discovery has deeply challenged the traditional sequence-structure-function paradigm, according to which protein function depends on a stable tertiary structure [8]. Actually, IDPs cover a spectrum of states that are in dynamic equilibrium under physiological conditions from fully structured, to molten globular, pre-molten globular, and random coil states [9].

Ever-increasing experimental evidence has revealed that IDPs are essential in many cellular processes [10-12] and plays critical roles in many human disease [13-16].

Sequences of IDPs are characterized by a marked bias in the amino acid composition, which are dramatically different from those of ordered proteins. The combination of a relatively low proportion of hydrophobic residues, a relatively high proportion of charged and polar residues, and high solvent accessibility represents an hallmark of IDPs [2,3,17]. All these features were shown to be associated with increased protein solubility [2,3,18], which is an important thermodynamic property in structural and biophysical studies.

In a recent work, we have defined an operational classification of the human proteome into three variants of proteins characterized by different functional spectra and roles in diseases [19]: i) ordered proteins (ORDPs), ii) structured proteins with intrinsically disordered regions (IDPRs), and iii) intrinsically disordered proteins (IDPs). We identified IDPs as those proteins having more than 30% of disordered residues (implying a shorter half-life [20]). We defined IDPRs as proteins with at least one long disordered segment (suggesting a higher sensitivity to proteolytic degradation [21]) but less than 30% of disordered residues in their sequence. Many studies aiming at a general classification of functional roles of protein intrinsic disorder were based on the required presence of long disordered regions in the sequence, thus considering IDPRs and IDPs as a single entity [10-12,22-25]. The main result of the study by Deiana et al. is that IDPs and IDPRs have different functional roles in the cell, and they deserve to be treated as distinct categories of proteins [19,26]. With this study, we emphasize this result by performing a systematic analysis of their amino acid compositions, their physical-chemical properties, and their structural aspects (both predicted and directly calculated on the structures in the Protein Data Bank). We concluded that IDPRs and IDPs are different, not only in terms of functional properties, roles in diseases, and evolutionary forces acting on them [19,26], but also as regards basic physical-chemical and structural properties.

## 2. Materials and Methods

### 2.1 Dataset sources

We downloaded the human proteome from the UniProt-SwissProt database (http://www.uniprot.org/uniprot) [27]. We selected human proteins by searching for reviewed proteins belonging to Homo Sapiens (Organism ID: 9606, Proteome ID: UP000005640). Human X-ray and NMR structures were retrieved from the Protein Data Bank (PDB) (https://www.rcsb.org) [28].

### 2.2 Disorder prediction

We identified disordered residues in the protein sequences from the UniProt-SwissProt database using MobiDB (http://mobidb.bio.unipd.it) [29], a consensus database that combines experimental data (especially from X-ray crystallography, nuclear magnetic resonance (NMR) and cryo-electron microscopy (Cryo-EM)), curated data and disorder predictions based on various methods. To predict protein disorder in pdb files, we used IUPred, which is a fast, robust, sequence-only predictor based on an energy estimation approach that captures the fundamental difference between the biophysical properties of ordered and disordered regions [30,31]. In particular, each residue in the sequence annotated in ‘SEQRES records’ in a given pdb file was classified as either ordered or disordered depending on whether the IUPred score was < 0.5 or > 0.5, respectively. Then, we used our criteria to partition the human proteins into variants (see next section).

### 2.3 Classification of disordered proteins in the human proteome

Following clear-cut criteria inspired by experimental observations [20,21], Deiana et al. partitioned the human proteome into three variants of disorder having different functional properties and roles in diseases: *i) ordered proteins* (ORDPs), *ii) proteins with intrinsically disordered regions* (IDPRs), *iii) intrinsically disordered proteins* (IDPs). Generally speaking, ORDPs are proteins with a limited number of disordered residues and the absence of disordered domains. IDPRs, unlike ORDPs, are proteins with at least one long disordered segment (implying a short half-life [21]) accommodated in globally folded structures. IDPs are proteins with a significant percentage of disorder (implying an unfolded structure and a high susceptibility to proteolytic degradation [20]).

### 2.4 Charge-Hydropathy plots

The Charge-Hydropathy (CH) plot is a tool to separate natively unfolded proteins from native folded ones [2]. In this plot, the mean net charge is plotted as the ordinate, and the mean hydrophobicity is plotted as the abscissa. The idea behind the CH-plot is that the combination of low mean hydrophobicity and high net charge let emerge natively unfolded conformations. Accordingly, it is possible to distinguish two different regions in the CH-plot associated with ordered and disordered proteins (here referred to as ordered and disordered phases, respectively). The separating line between ordered and disordered phases we used here is the original one by Uversky et al. [2]:

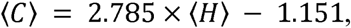

where ⟨*C*⟩ and ⟨*H*⟩ are the mean net charge and the mean hydrophobicity of the protein, respectively. The mean hydrophobicity was estimated using the Kyte-Doolittle normalized scale [32].

### 2.5 Mean packing scale

To estimate the mean packing of a protein from its sequence without relying on the structure, we used the packing scale by Galzitskaya et al. [33]. Considering a training set of 6626 protein structures extracted from the SCOP database, they estimated the average number of close residues for each of 20 amino acids. Using these data, the protocol for estimating the mean packing of a protein is the following. We assign to each residue in the sequence under study one of the 20 values according to the scale by Galzitskaya et al. [33]. The position-specific estimations of packing were averaged over a window of 21 residues, and the average value was assigned to the central residue of the window (taking into account the limitations on both sides of the protein sequence).

### 2.6 Mean pairwise energy scale

To estimate the structural stability of a protein from its sequence without relying on the structure, we used the energy estimation approach at the core of the IUPred disorder prediction method [30,31]. The key component of the calculations is the energy estimation matrix, a 20 by 20 matrix, whose elements characterize the general preference of each pair of amino acids to be in contact as derived from globular proteins. For the actual estimation of the pairwise energy for each residue, the position-specific estimations of energies were averaged over a window of 21 residues, and the average value was assigned to the central residue of the window. Then, the average energy per residue (here referred to as ‘mean pairwise energy’) is approximated by the arithmetic mean of the pairwise energy values of all residues that compose the protein sequence.

### 2.7 Theoretical approach to the radius of gyration

In polymer physics and structural biology, the radius of gyration is an important physical quantity, which is often used to describe the dimensions and compactness of a polymer. For a given protein configuration taken from the PDB, we compute the radius of gyration as follows:

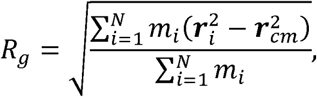

where 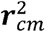 is the position of the center of mass of the protein, 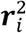 is the position of atom *i*, and *m*_*i*_ is its mass. In the Flory theory, the radius of gyration of a homopolymer at the equilibrium state depends on the chain length through a scaling law *R*_*g*_ ∼ *N*^*ν*^, where *ν* is referred to as ‘Flory exponent’ and depends on the solvent conditions [34,35]. In poor solvent conditions, the polymer chain is strongly compressed by the pressure exerted by the solvent, and the excluded volume effect 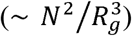 and the three-body repulsive interaction 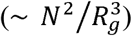 constitute the main part of the total energy. Thus, minimizing this energy with respect to *R*_*g*_ gives the 1/3 law. On the other hand, in good solvent conditions, the polymer chain is extended, and the excluded volume effect 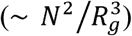 and the entropy effect 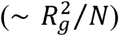 become the dominant terms in the energy, thus implying the 3/5 law. However, proteins are heteropolymers made up of 20 different amino acids, with different physical chemical properties (e.g., hydrophobicity, charge). For instance, if all residues are hydrophobic, which is equivalent to poor solvent conditions, the protein will be highly compressed by solvent pressure and the scaling exponent will be *ν* ∼ 1/3. On the other hand, if all residues are hydrophilic, which is equivalent to good solvent conditions, the protein will be extended and the scaling exponent will be *ν* ∼ 3/5. Thus, for a general protein under physiological conditions, which can contain both hydrophobic or hydrophilic residues, the scaling exponent will be intermediate between 1/3 and 3/5.

Using the generalized formula for the radius of gyration by Hong and Lei [35], it is simple to demonstrate that

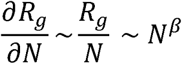

So studying the ratio *R*_*g*_/*N* as a function of *N*, it is possible to infer important information on *∂R*_*g*_/*∂N* such as the scaling exponent *β*, which gives an idea about how the radius of gyration varies by adding a monomers to protein.

### 2.8 Prediction and analysis of protein solubility

Protein solubility was assessed using Protein-Sol (https://protein-sol.manchester.ac.uk) [36], which is reasonably accurate, fast, and available for local use. It processes amino acid sequence and calculates predicted solubility. Thirty-five features are considered in the algorithm, 20 amino acid compositions; 7 composites: K-R, D-E, K+R, D+E, K+R-D-E, K+R+D+E, F+W+Y; and 8 further predicted features: length, pI, hydropathy, absolute charge at pH 7, fold propensity, disorder, sequence entropy, and β-strand propensity.

### 2.9 Analyses of the accessible surface area

Accessible surface area (ASA), i.e., the surface area of a biomolecule that is accessible to a solvent, was calculated using STRIDE [37]. STRIDE is a software available for local use that takes a single PDB file as an input and predicts the ASA of each residue in the structure. Then, an average ASA was assigned to each protein in the pdb dataset as the arithmetic mean of the ASA values of its residues.

## 3 RESULTS

### 3.1 Amino acid compositions of the variants

The amino acid composition of a protein strongly influences its capacity to fold in a well-stable structure. In line with this, the three variants of disorder are differentially enriched in order- and disorder-promoting amino acids (Fig. 1). ORDPs are enriched in order-promoting amino acids and depleted in disorder-promoting amino acids; conversely, IDPs are enriched in disorder-promoting amino acids and depleted in order-promoting amino acids. IDPRs show a less clear pattern of amino acid usage with a slight preference for order-promoting amino acid, thus suggesting the presence of a well-folded structure in these proteins. In support of this, IDPRs are more similar to ORDPs than to IDPs in terms of amino acid composition (see clustering on the bottom panel in Fig. 1).

**Figure 1.**
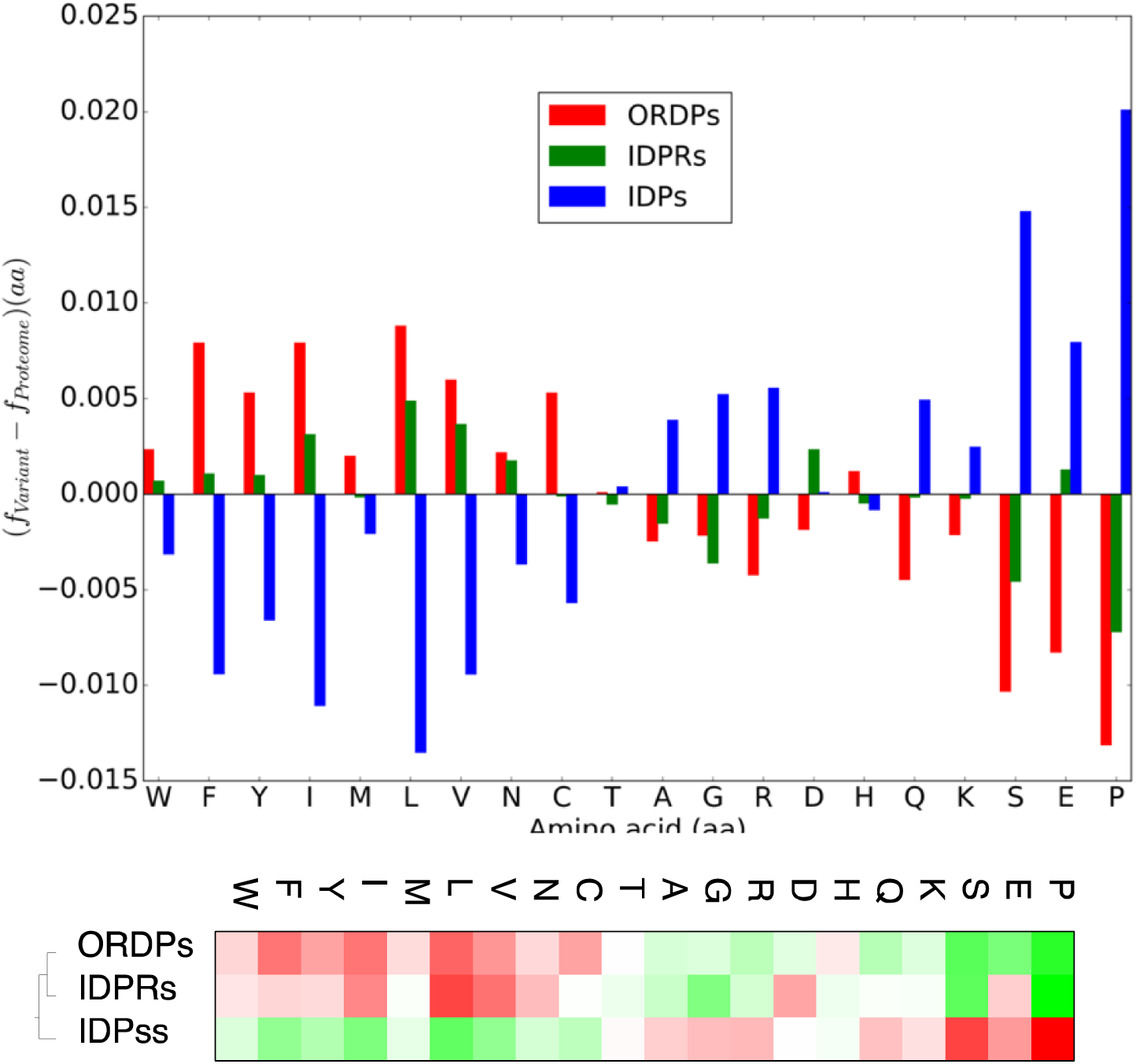
Differential amino acid compositions of ORDPs, IDPRs, IDPs, with respect to the human proteome. On the top, the bar charts of the normalized differential amino acid compositions of ORDPs (in red), IDPRs (in green), and IDPs (in blue), with respect to the overall amino acid composition of the human proteome (the reference). The 20 amino acids are abbreviated according to standard convention and ordinated from order-to disorder-promoting [38]. On the bottom, clustering of ORDPs, IDPRs, and IDPs based on the normalized differential amino acid compositions shown in the top panel. The clustering was obtained through the CLUTO toolkit: (http://glaros.dtc.umn.edu/gkhome/cluto/cluto/overview).

### 3.2 Charge-hydropathy plots of the variants

We use the charge-hydropathy (CH) plot to characterize ORDPs, IDPRs, and IDPs in terms of the net charge and mean hydrophobicity, two crucial parameters in determining the capacity of a protein to fold (Fig. 2). Coherently with their enrichment in order-promoting amino acid, ORDPs and IDPRs are preferentially located in the ordered phase, as expected for well-structured proteins. On the contrary, IDPs are widespread on both sides of the separating line with a preference towards the disordered phase, as expected for unstructured proteins. This is clearer by looking at the centroids of the protein variants: both the centroids of ORDPs and IDPRs are located in the ordered phase of the CH-plot, whereas the centroid of IDPs is near the border line with a larger overlap with the disordered phase. In line with Gsponer et al. [20], a percentage of disordered residues equal to 30% appears to be more proper for discriminating ordered proteins from disordered ones. In particular, if the percentage of disordered residues is less than 30% then the centroid is in the ordered phase, whereas increasing the percentage of disordered residues more than 30%, the centroids of IDPs cross the separating line moving towards the disordered phase.

**Figure 2:**
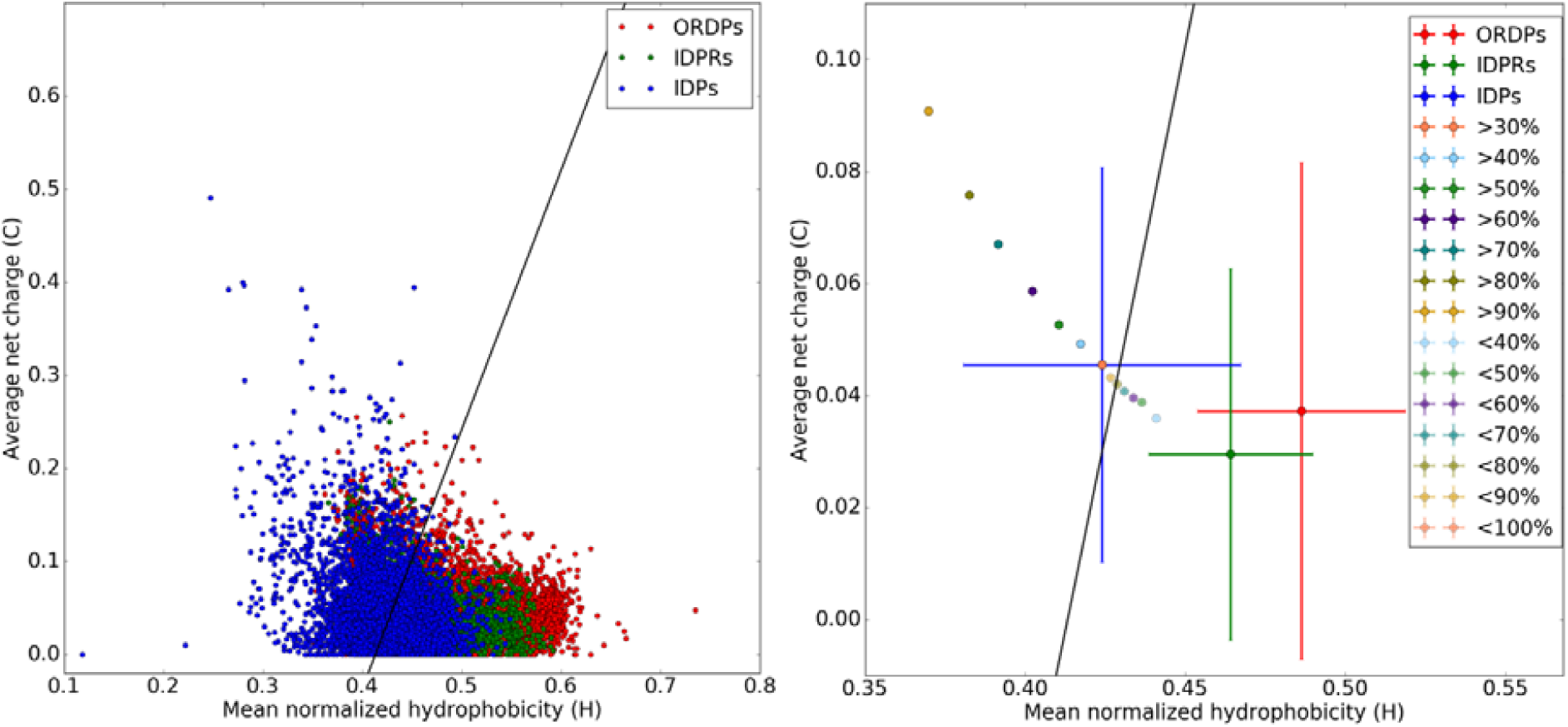
Charge-hydropathy plot of ORDPs, IDPRs, and IDPs. On the left, the net charge is plotted as the ordinate and the mean hydrophobicity is plotted as the abscissa for ORDPs (red), IDPRs (green), and IDPs (blue). The solid black line represents the border between ordered and disordered phases in the CH-plot:. On the right, the centroids of ORDPs (red), IDPRs (green), IDPs (blue), and for different percentages of disordered residues (more than a given threshold, that is >30%, >40%, and so on, or up to a given threshold, that is <40%, <50%, and so on). Errors bars are the standard deviations of distributions.

### 3.3 Representation of the variants in the MP-MPE plane

To structurally characterize our variants of disorder, we introduce the *mean packing vs*. *mean pairwise energy (MP-MPE) plane*. In this plot, the mean packing value is plotted as the ordinate, and the mean pairwise energy value is plotted as the abscissa. In Fig. 3, we report the MP-MPE plane for the human proteome, separated into the three variants.

**Figure 3.**
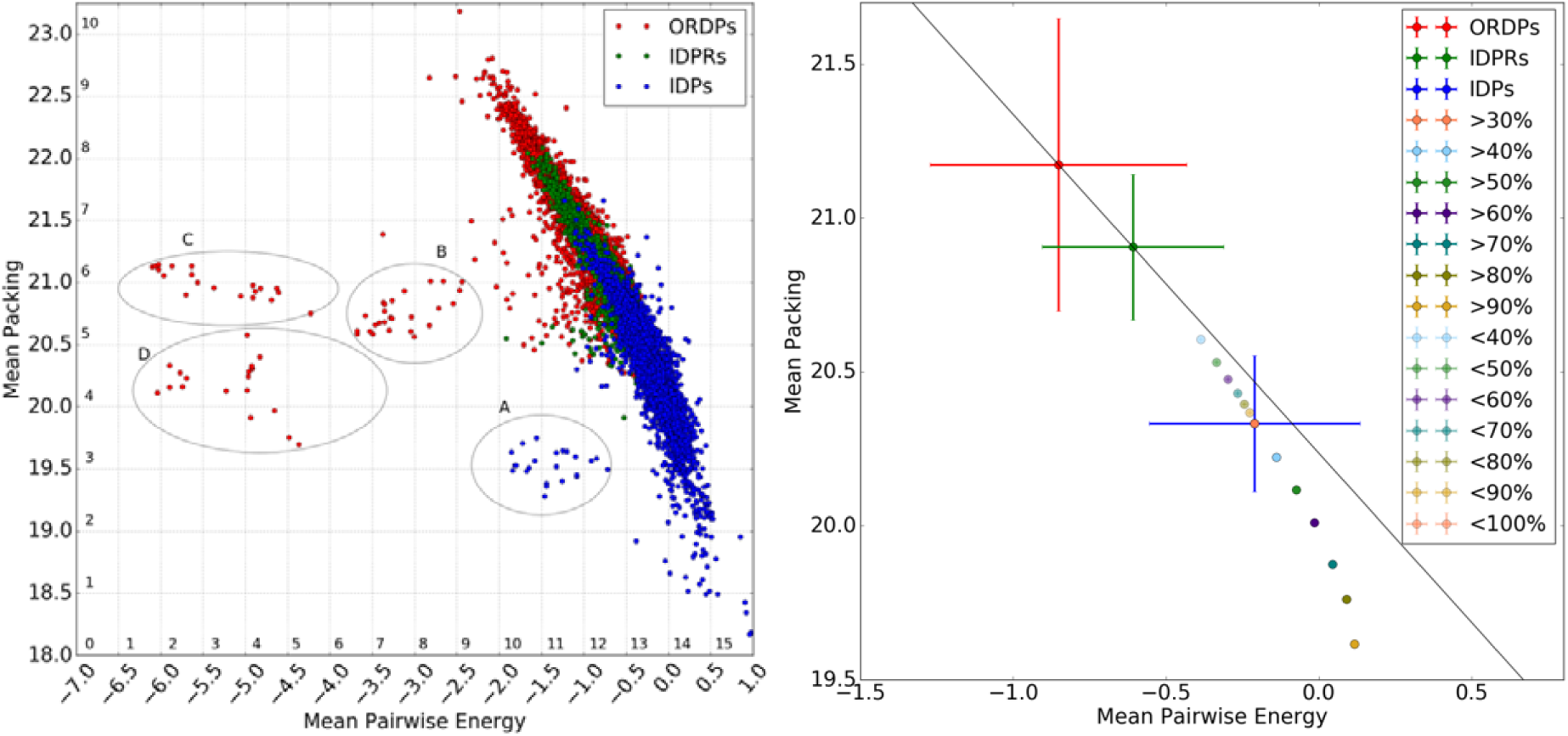
MP-MPE plane of ORDPs, IDPRs, and IDPs. On the left, MP-MPE plane for the human proteome, taken here as a case of study. Each point represents a single protein. ORDPs, IDPRs, and IDPs are reported in red, green, and blue, respectively. As expected, MP and MPE are strongly correlated because the higher is the packing, the higher is the expected number of inter-residue interactions. The solid line in both graphs is the regression line only considering non-disordered variants (ORDPs and IDPRs) () that we refer to as the ‘principal sequence’. ORDPs, IDPRs, and IDPs are in this order along the principal sequence. Increasing the percentage of disorder, there is a systematic sliding along the principal sequence. Notably, increasing the percentage of disordered residues more than 30% (threshold that we used to classify IDPs), proteins start to deviate from the principal sequence. On the right, the centroids of ORDPs, IDPRs, and IDPs in the MP-MPE plane. The error bars denote the standard deviation of distribution of points around the centroids. Colored circles without uncertainty bars are the centroids of IDPs with different percentages of disordered residues (>30%, >40%, >50%, >60%, >70%, >80%, >90%, or <40%, <50%, <60%, <70%, <80%, <90%, <100%). Four clusters of outlier proteins that strongly break the MP-MPE correlation are denoted as: A, B, C, and D.

A strong negative correlation is observed between MP and MPE because the more packed is the structure, the higher is the propensity to make inter-residue contacts. In line with what observed in CH-plot, the percentage of disordered residues acts as a control parameter along the regression line (here referred to as *principal sequence*). In particular, the higher is the disorder content of a protein, the lower is its position on the principal sequence. Looking at the centroids in Fig. 3, ORDPs, IDPRs, and IDPs are in this order along the principal sequence, each one in a typical range of MP and MPE. ORDPs occupy the top-right region, indicating that these proteins have a high propensity to fold into compact and stable structures. IDPRs are located in the central region, having intermediate MP and MPE values between ORDPs and IDPs. Finally, IDPs are located in the bottom-left region of the principal sequence, being less compact and more flexible than ORDPs and IDPRs. Interestingly, increasing the percentage of disordered residues more than 30%, proteins deviate from the principal sequence, breaking the linear correlation between MP and MPE (Fig. 3). It is worth noting the presence of four clusters, characterized by much smaller MPE values than the rest of human proteins (Fig. 3). All these outlier proteins are mostly related to keratin proteins and they are very rich in cysteine residues (Fig. 4), which stabilize their structure through the formation of disulfide bridges (see Table S1, S2, S3, S4 for the UniProt ID and a brief description of these proteins). In line with that, the departure from the principal sequence is linearly correlated with the fraction of cysteines in each cluster, i.e., the higher is the content of cysteines in the cluster, the greater is the distance of such cluster from the principal sequence.

**Figure 4.**
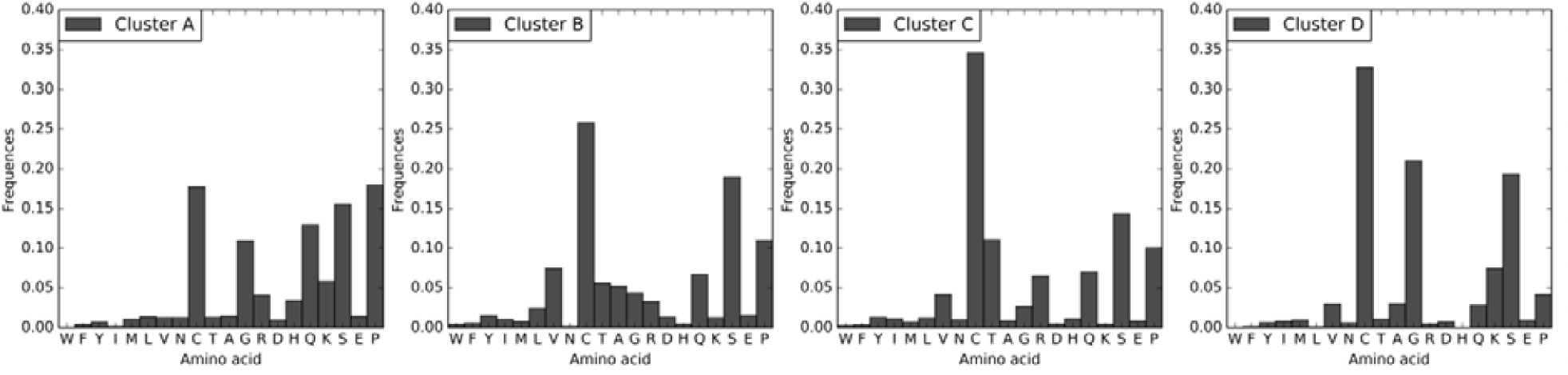
Amino acid compositions of the four clusters (A, B, C, D) of outlier proteins. The 20 amino acids are abbreviated according to standard convention and ordinated from order-to disorder-promoting [38]. The most representative residue is the cysteine (C).

### 3.4 The structures of ORDPs, IDPRs, and IDPs are characterized by different scaling laws

Based on Flory’s polymer theory, we have characterized our variants of disorder in terms of the exponent (), which describes how the radius of gyration scales with the chain length and gives an idea of the compactness of the protein. In Fig. 5, we report the log-log plot of (measured for each protein structure) as a function of the number of residues that compose the proteins. Coherently with the results by [35], we found a scaling exponent for all variants of disorder, indicating that all proteins in our variants are more compact than polymers in good solvent, but looser than highly compressed polymers in poor solvent. In particular, Flory exponent increases with the percentage of disordered residues that characterizes the variant (see insert in Fig. 5). Thus, IDPs have the highest Flory exponent, thus implying a more extended conformations than ORDPs and IDPRs; conversely, ORDPs have the lowest Flory exponent, underscoring their globular characteristics. IDPRs have an intermediate Flory exponent between ORDPs and IDPs, likely due to the presence of long disordered segments that swells their structures.

Next, we studied how the addiction of a single monomer influences the radius of gyration of the protein, given the fact that such protein belongs to one of the variants. As shown in Materials and Methods, to perform this analysis, it is enough to study the trend of as a function of. In Fig. 6, we report measured for each protein structure as a function of the number of residues that compose the proteins.

**Figure 5:**
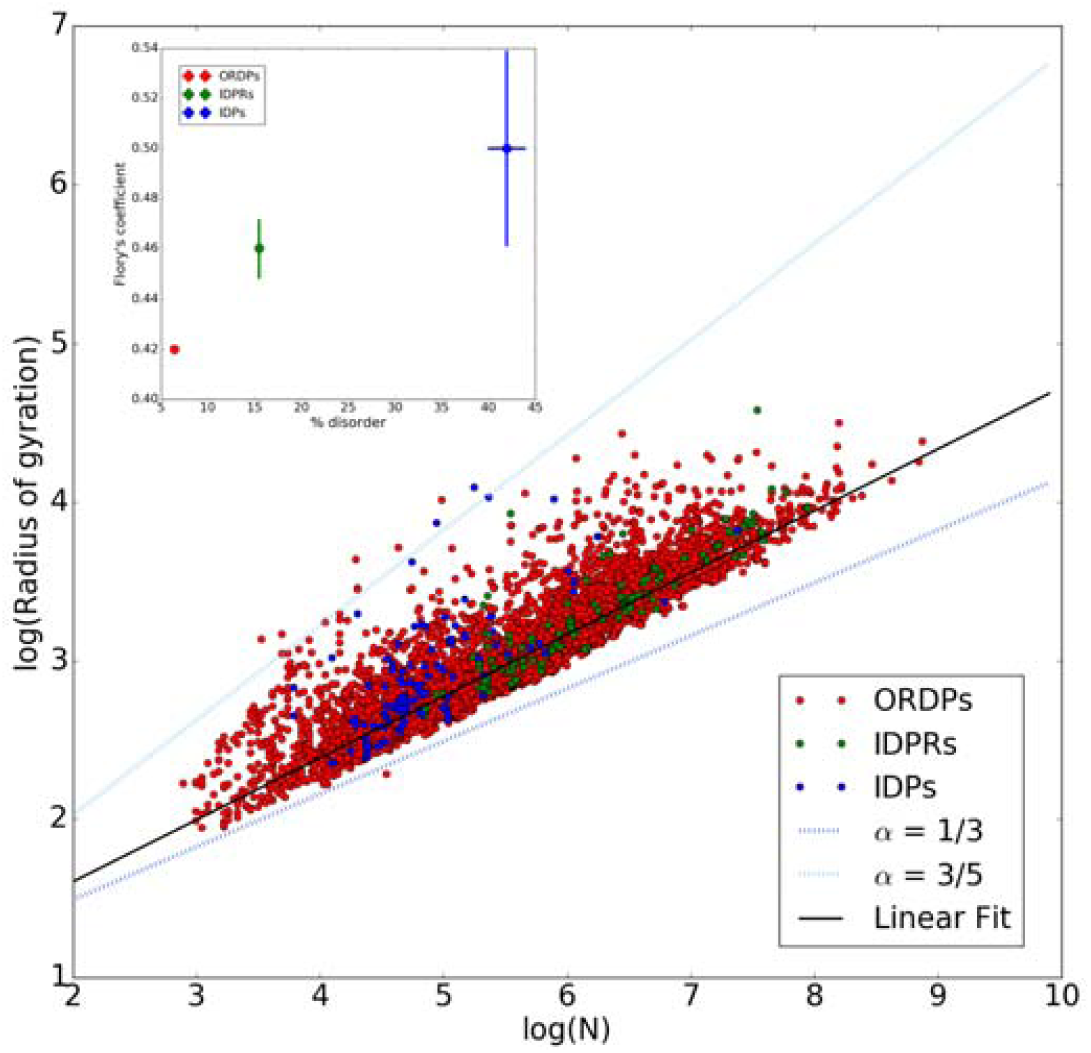
Log-log plot of the radius of gyration and protein length for ORDPs (in red), IDPRs (in green), and IDPs (in blue). The dotted lines represent the special case predicted in the Flory theory for poor (in blue) and good (in cyan) solvent conditions. The solid black line represents the linear fit of the whole human proteome. In the insert, we report the Flory coefficient as a function of the average percentage of disordered residues for the three variants of proteins. The error bars denote the standard errors of the linear regressions.

**Figure 6:**
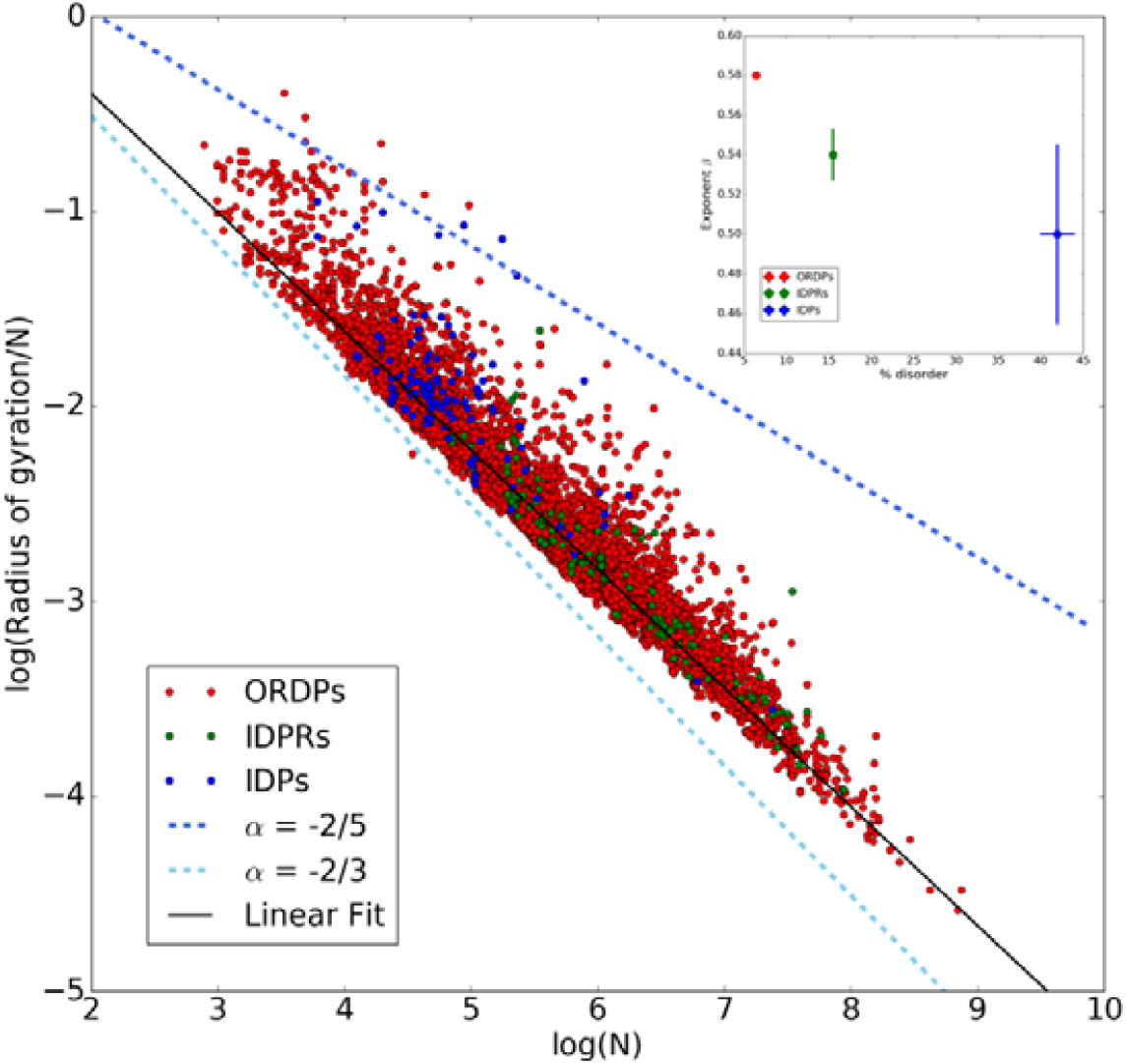
Log-log plot of the ratio and protein length for ORDPs (in red), IDPRs (in green), and IDPs (in blue). The dotted lines represent the special case predicted for poor (in blue) and good (in cyan) solvent conditions. The solid black line represents the linear fit of the whole human proteome. In the insert, we report the scaling exponent as a function of the average percentage of disordered residues for the three variants of proteins. The error bars denote the standard errors of the linear regressions.

All variants of human proteins have a scaling exponent *β* which is intermediate between the cases of poor and good solvent conditions (i.e., −2/5<*β*<−2/3). As expected, *β* decreases with the percentage of disordered residues, because more packed is the protein structure, less is the space to accommodate a new monomer. Indeed, each monomer added to a well-packed protein will be positioned almost certainly on the external surface, leading to a faster growth of the radius of gyration. Thus, IDPs and ORDPs have the lowest and the highest scaling exponent *β*, respectively. IDPRs have a scaling exponent which is intermediate between ORDPs and IDPs, likely due to the presence of a long disordered segments in their structures. At this point a remark is in order. To measure *R*_*g*_, as we did above, one needs to have a protein structure from the PDB. Clearly, attributing a gyration radius to an IDP should be an oxymoron. Indeed it is well known that the protein Data Bank contains a considerable amount of predicted disorder, this is not a scandal, but see references herein [39,40].

### 3.5 IDPs are more soluble and have higher solvent accessibility than ORDPs and IDPRs

Protein solubility and solvent accessibility are two important features that distinguish ordered proteins from disordered ones. Ordered proteins are typically constituted of an internal core (composed by bulky hydrophobic amino acids) and an external surface, which is usually enriched in polar and charged amino acids that interact favorably with water, leading to aqueous solubility. The presence of hydrophobic surface residues (e.g., binding sites for other proteins) and the denaturation of otherwise folded proteins lead to the exposure of hydrophobic residues to water and reduce solubility, sometimes leading to aggregation, precipitation, and disease [21]. On the other hand, intrinsically disordered proteins lack of a well-defined structure and have a more relaxed conformation because their sequences are depleted in hydrophobic amino acids. The accompanying enrichment in polar and charged amino acids, as a general rule, causes intrinsically disordered proteins to be soluble in aqueous solutions [18,21] and to have higher solvent accessibility than globular proteins [41-43]. Coherently, we found that IDPs are characterized by the highest values of solubility and of accessible surface area with respect to the rest of the human proteome (Fig. 7). Conversely, ORDPs and IDPRs have comparable solubilities, likely due to their similar structural, globular properties.

**Figure 7:**
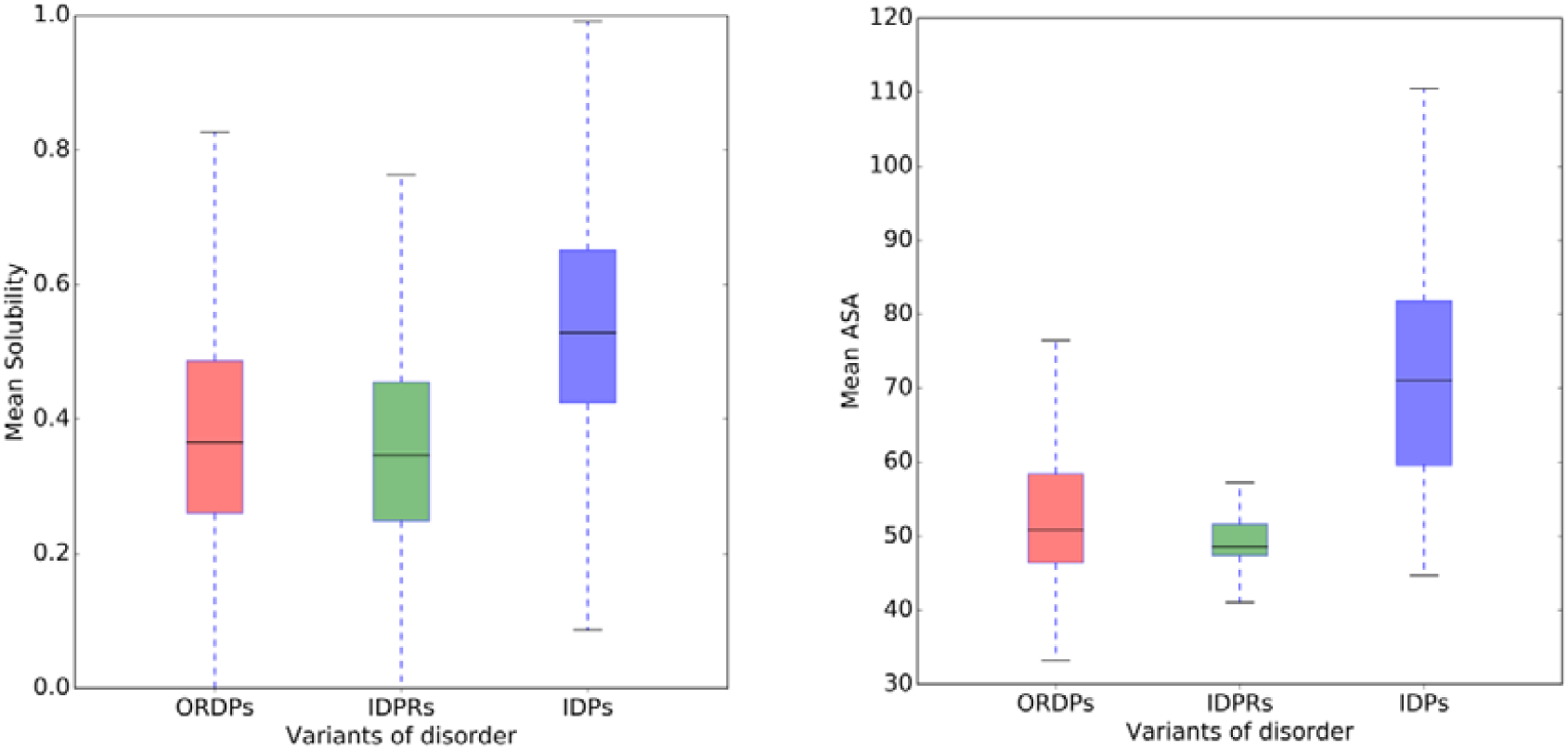
Solubility and Accessible Surface Area. Box plots of protein solubilities (on the left) and accessible surface areas (on the right) of ORDPs, IDPRs, and IDPs. IDPs are significantly characterized by the highest values of solubility and accessible surface area with respect to the rest of human proteome (Kruskal-Wallis H-test and Mann-Whitney U-test with Bonferroni correction for multiple comparisons, p-value<0.00001).

## 4. Discussion

In a previous study, we introduced an operationally classification of the human proteome in three variants of disorder having different functional spectra and roles in diseases [19]: i) ordered proteins (ORDPs), ii) structured proteins with intrinsically disordered protein regions (IDPRs), and iii) intrinsically disordered proteins (IDPs). That classification was effective in functionally separating IDPRs from IDPs, which up until now have been generally considered as a whole [10-12,22-25]. Furthermore, performing systematic analyses of these variants in terms of evolutionary forces acting on them, we have recently shown that IDPRs on the one side are subject to different patterns of evolutionary pressures with respect to IDPs and ORDPs on the other one [26].

The main outcome of the present study, in a nutshell, is that IDPRs have a structural signature more similar to that of ORDPs than to IDPs.

Firstly, we focused on the amino acid compositions. ORDPs are enriched in order-promoting amino acid and depleted in disorder-promoting ones. Conversely, IDPs are enriched in disorder-promoting amino acid and depleted in order-promoting ones. Both these observations are coherent with the concept of ORDPs as opposed to IDPs. As regards to IDPRs, their amino acid composition is intermediate between those of ORDPs and IDPs, with a slight propensity for order-promoting amino acids, thus suggesting a general picture of this variant as folded proteins accommodating significant disorder regions in their structure. IDPRs appear to be heterogeneous “mosaic of units” encompassing patterns of of *foldons* (i.e. ordered segments), *semi-foldons* (i.e., partially folded protein segments with transient residual structure), *inducible foldons* (i.e., segments that are able to fold at interaction with specific binding partners), *nonfoldons* (i.e., protein segments with functions depending on their intrinsically disordered nature), and *unfoldons* (i.e., segments that are able to unfold under specific conditions, becoming functional) [9,44].

According to their preference for order-promoting amino acids, we find that both ORDPs and IDPRs are preferentially located in the ordered charge-hydropathy phase where mean hydrophobicity and mean net charge are typical of well-structured proteins. Conversely, IDPs are preferentially located in the disordered phase, where the combination of low mean hydrophobicity and relatively high net charge support their being structureless as the true natively unfolded proteins. As expected, increasing the percentage of disordered residues over 30%, IDPs move away from the separating line of the CH-plot, towards the disordered phase.

Next, we introduced the Mean Packing – Mean Pairwise Energy (MP-MPE) plane, which allowed us to structurally characterize (ab-initio, sequence-only) ORDPs, IDPRs, and IDPs, even in the absence of a structural model. A strong negative correlation between MP and MPE is observed with the emergence of a ‘principal sequence’ where most of the proteins are located. The percentage of disordered residues acts as a parameter controlling the sliding of proteins along the principal sequence. ORDPs, i.e., well-structured proteins, are located on the top-left of the plane, in the region corresponding to high MP values and low MPE values. To follow, we find IDPRs with slightly lower MP values and slightly higher MPE values than ORDPs. IDPs are located at the bottom, in the region corresponding to lower MP values and higher MPE values. Interestingly, ORDPs and IDPRs appear to be characterized by a strong linear correlation between their values of MP and MPE. This is in line with what we expect for well-structured globular proteins: the higher is the packing, i.e., the expected average number of close residues, the higher is the expected number of contacts that stabilize the native state. On the contrary, IDPs do not satisfy rigidly the proportionality between MP and MPE with a gradual divergence from the linear relation that characterize ORDPs and IDPRs. In particular, the more is the percentage of disordered residues in IDPs, the larger is their deviations from the principal sequence. We suppose that this peculiar feature of IDPs is probably due to the fact that energetic interactions in these proteins are apparently insufficient for compensating for the conformational entropy, whose contribution, by the way, is not considered in the mean pairwise energy calculation. Thus, we conclude that the departure from the principal sequence results from a shift in the structural properties, possibly dominated, as often stressed, by entropic effects that are also mirrored in the low hydrophobicity and high net charge that characterize IDPs in the CH-plot.

We also studied our variants of disorder in terms of scaling exponents of the radius of gyration with the length of human proteins that we can retrieve in the Protein Data Bank. Our results show that IDPs have the highest Flory exponent, thus confirming that IDPs have a more extended ensemble of conformation than ORDPs and IDPRs. With a similar reasoning, based on the scaling exponent of the first derivative of the radius of gyration, we analyzed how the addition of a single monomer increases the radius of gyration of the protein, in each of the variants. Reasonably, IDPs are more flexible than ORDPs and IDPRs in increasing their length without increasing their radius of gyration (i.e., IDPs have a lower scaling exponent *β*).

Finally, we performed an analysis of protein solubility and accessible surface area, two interdependent parameters both related to interfacial interactions with the solvent. We found that IDPs are characterized by the highest values of solubility and of accessible surface area with respect to the rest of the human proteome. Generally speaking, this is clear considering that, in general, proteins with more charged and polar residues on their accessible surface to water molecules show higher solubility [45].

## 5. Conclusions

In summary, our results corroborate our previous claim to consider IDPRs and IDPs as distinct protein variants [19,26]. At the same time, we identify IDPs as the true natively unfolded proteins, having peculiar functional and evolutionary features but also distinct physical-chemical and structural properties. As a final remark, let us observe that the keratin-related human proteins that clusterize out of the principal sequence, in the MP-MPE plane, are quite relevant in an evolutionary perspective. Indeed, a recent paper has shown as hairs, feathers, and scales of present-day species evolved from an ancestral anatomical placode [46]. It would be of interest to investigate the degree of conservation of these outliers, in the MP-MPE plane.

## Supporting information

see Table S1, S2, S3, S4 for the UniProt ID and a brief description of these proteins

## Abbreviations

ORDPs: ordered proteins
IDPRs: structured proteins with intrinsically disordered regions
IDPs: intrinsically disordered proteins.

