## Supplementary material for "Variants of intrinsic disorder: structural characterization": see Table S1, S2, S3, S4 for the UniProt ID and a brief description of these proteins

**Supplemetary information**

| **CLUSTER A** | |
| --- | --- |
| **Gene ID** | **Protein** |
| Q5TCM9 | Late cornified envelope protein 5A |
| P22532 | Small proline-rich protein 2D |
| Q9BYE3 | Late cornified envelope protein 3D |
| Q9BYE4 | Small proline-rich protein 2G |
| P35326 | Small proline-rich protein 2A |
| Q5T5A8 | Late cornified envelope protein 3C |
| Q5TA81 | Late cornified envelope protein 2C |
| Q5TA82 | Late cornified envelope protein 2D |
| Q5T751 | Late cornified envelope protein 1C |
| Q5T752 | Late cornified envelope protein 1D |
| Q5T753 | Late cornified envelope protein 1E |
| Q5T754 | Late cornified envelope protein 1F |
| Q9UGL9 | Cysteine-rich C-terminal protein 1 |
| P35325 | Small proline-rich protein 2B |
| Q5T7P3 | Late cornified envelope protein 1B |
| Q5T7P2 | Late cornified envelope protein 1A |
| Q156A1 | Ataxin-8 |
| Q5TA78 | Late cornified envelope protein 4A |
| Q5TA79 | Late cornified envelope protein 2A |
| Q5TA76 | Late cornified envelope protein 3A |
| Q5TA77 | Late cornified envelope protein 3B |
| Q5T871 | Late cornified envelope-like proline-rich protein 1 |
| Q96RM1 | Small proline-rich protein 2F |
| P22531 | Small proline-rich protein 2E |
| P49901 | Sperm mitochondrial-associated cysteine-rich protein |
| Q5T5B0 | Late cornified envelope protein 3E |

**Table S1: List of outlier proteins in the cluster A in Fig. 3.**

| **CLUSTER B** | |
| --- | --- |
| **Gene ID** | **Protein** |
| Q9BYR9 | Keratin-associated protein 2-4 |
| P60369 | Keratin-associated protein 10-3 |
| P60368 | Keratin-associated protein 10-2 |
| P0C7H8 | Keratin-associated protein 2-3 |
| Q9BYT5 | Keratin-associated protein 2-2 |
| P60014 | Keratin-associated protein 10-10 |
| P60409 | Keratin-associated protein 10-7 |
| Q07627 | Keratin-associated protein 1-1 |
| Q8IUG1 | Keratin-associated protein 1-3 |
| P59990 | Keratin-associated protein 12-1 |
| P59991 | Keratin-associated protein 12-2 |
| P60331 | Keratin-associated protein 10-1 |
| P60413 | Keratin-associated protein 10-12 |
| P60412 | Keratin-associated protein 10-11 |
| P60411 | Keratin-associated protein 10-9 |
| P60410 | Keratin-associated protein 10-8 |
| Q9BYU5 | Keratin-associated protein 2-1 |
| Q3LI58 | Keratin-associated protein 21-1 |
| Q3LI59 | Keratin-associated protein 21-2 |
| Q9BYS1 | Keratin-associated protein 1-5 |
| P60372 | Keratin-associated protein 10-4 |
| P60370 | Keratin-associated protein 10-5 |
| P60371 | Keratin-associated protein 10-6 |
| P0C5Y4 | Keratin-associated protein 1-4 |

**Table S2: List of outlier proteins in the cluster B in Fig. 3.**

| **CLUSTER C** | |
| --- | --- |
| **Gene ID** | **Protein** |
| A8MVA2 | Keratin-associated protein 9-6 |
| Q9BYP9 | Keratin-associated protein 9-9 |
| Q9BYR0 | Keratin-associated protein 4-7 |
| Q9BYR3 | Keratin-associated protein 4-4 |
| Q9BYR5 | Keratin-associated protein 4-2 |
| Q9BYR4 | Keratin-associated protein 4-3 |
| A8MXZ3 | Keratin-associated protein 9-1 |
| Q9BYQ4 | Keratin-associated protein 9-2 |
| Q9BYQ5 | Keratin-associated protein 4-6 |
| Q9BYQ6 | Keratin-associated protein 4-11 |
| Q9BYQ7 | Keratin-associated protein 4-1 |
| Q9BYQ0 | Keratin-associated protein 9-8 |
| Q9BYQ2 | Keratin-associated protein 9-4 |
| Q9BYQ3 | Keratin-associated protein 9-3 |
| A8MTY7 | Keratin-associated protein 9-7 |
| Q9BQ66 | Keratin-associated protein 4-12 |
| Q9BYQ8 | Keratin-associated protein 4-9 |
| Q9BYQ9 | Keratin-associated protein 4-8 |

**Table S3: List of outlier proteins in the cluster C in Fig. 3.**

| **CLUSTER D** | |
| --- | --- |
| **Gene ID** | **Protein** |
| P80297 | Metallothionein-1X |
| P80294 | Metallothionein-1H |
| P04731 | Metallothionein-1A |
| Q93083 | Metallothionein-1L |
| P0DM35 | Metallothionein1H-like protein 1 |
| P26371 | Keratin-associated protein 5-9 |
| P02795 | Metallothionein-2 |
| Q6L8G8 | Keratin-associated protein 5-7 |
| Q6L8G9 | Keratin-associated protein 5-6 |
| P25713 | Metallothionein-3 |
| P04732 | Metallothionein-1E |
| Q6L8G4 | Keratin-associated protein 5-11 |
| Q6L8G5 | Keratin-associated protein 5-10 |
| P04733 | Metallothionein-1F |
| Q6L8H4 | Keratin-associated protein 5-1 |
| P13640 | Metallothionein-1G |
| Q6L8H2 | Keratin-associated protein 5-3 |
| Q6L8H1 | Keratin-associated protein 5-4 |
| Q701N4 | Keratin-associated protein 5-2 |
| O75690 | Keratin-associated protein 5-8 |
| Q701N2 | Keratin-associated protein 5-5 |
| P07438 | Metallothionein-1B |
| A1L3X4 | Putative metallothionein MT1DP |

**Table S4: List of outlier proteins in the cluster D in Fig. 3.**
